# Phylogenomics provides new insights into gains and losses of selenoproteins among Archaeplastida

**DOI:** 10.1101/674895

**Authors:** Hongping Liang, Tong Wei, Yan Xu, Linzhou Li, Sunil Kumar Sahu, Hongli Wang, Haoyuan Li, Xian Fu, Gengyun Zhang, Michael Melkonian, Xin Liu, Sibo Wang, Huan Liu

**Author notes:** these authors contributed equally to this work (Hongping Liang, Tong Wei, Yan Xu).

## Abstract

Selenoproteins that contain selenocysteine (Sec) are found in all kingdoms of life. Although they constitute a small proportion of the proteome, selenoproteins play essential roles in many organisms. In photosynthetic eukaryotes, selenoproteins have been found in algae but are missing in land plants (embryophytes). In this study, we explored the evolutionary dynamics of Sec incorporation by conveying a genomic search for the Sec machinery and selenoproteins across Archaeplastida. We identified a complete Sec machinery and variable sizes of selenoproteomes in the main algal lineages. However, the entire Sec machinery was missing in the BV clade (Bangiophyceae-Florideophyceae) of Rhodoplantae (red algae) and only partial machinery was found in three species of Archaeplastida, indicating parallel loss of Sec incorporation in different groups of algae. Further analysis of genome and transcriptome data suggests that all major lineages of streptophyte algae display a complete Sec machinery, although the number of selenoproteins is low in this group, especially in subaerial taxa. We conclude that selenoproteins tend to be lost in Archaeplastida upon adaptation to a subaerial or acidic environment. The high number of redox-active selenoproteins found in some bloom-forming marine microalgae may be related to defense against viral infections. Some of the selenoproteins in these organisms may have been gained by horizontal gene transfer from bacteria.

## 1. Introduction

Selenium (Se) is an essential trace element for human health and its deficiency leads to various diseases, such as Keshan and Kashin-Beck diseases, and affects the immune system and promotes cancer development [1,2]. An essential Se metabolism is present in many organisms, including bacteria, archaea, and eukaryotes [1,3,4]. However, higher concentrations of Se are toxic by functioning as a pro-oxidant, which affects the intracellular glutathione (GSH) pool leading to an enhanced level of ROS accumulation [5,6]. Se is essential for growth and development of numerous algal species but not for terrestrial plants (embryophytes), although it accumulates in certain plant species and can serve as dietary sources for Se uptake [3,7-9].

Se is incorporated into nascent polypeptides in the form of selenocysteine (Sec), the 21st amino acid [10]. Se incorporation requires a specialized machinery and Sec insertion sequence (SECIS) elements present in selenoprotein mRNAs [11,12]. In eukaryotes, it consists of Sec synthesis and Sec incorporation. Sec synthesis starts with tRNA^Sec^, aminoacylated with serine, which is phosphorylated by O-phosphoseryl-transfer tRNA^Sec^ kinase (PSTK) and then catalyzed by Sec synthase (SecS) to produce selenocysteinyl-tRNA^Sec^ from selenophosphate [10-13]. The Sec donor, selenophosphate, is generated from selenide by selenophosphate synthetase 2 (SPS2), which is often a selenoprotein itself [14,15]. During Sec incorporation, SECIS-binding protein 2 (SBP2) recognizes the SECIS elements in the 3’-untranslated region (3’-UTR) and recruits the Sec-specific elongation factor (eEFSec) that delivers selenocysteinyl-tRNA^Sec^ to the ribosome at the in-frame Sec-coding UGA stop codon (Figure 2a). Bacteria possess a similar machinery including *selB* (Sec-specific elongation factor), *selC* (tRNA^Sec^) and *selD* (selenophosphate synthase), except that Sec synthesis is catalyzed by a single bacterial Sec synthase, *SelA* [10].

**Figure 1.**
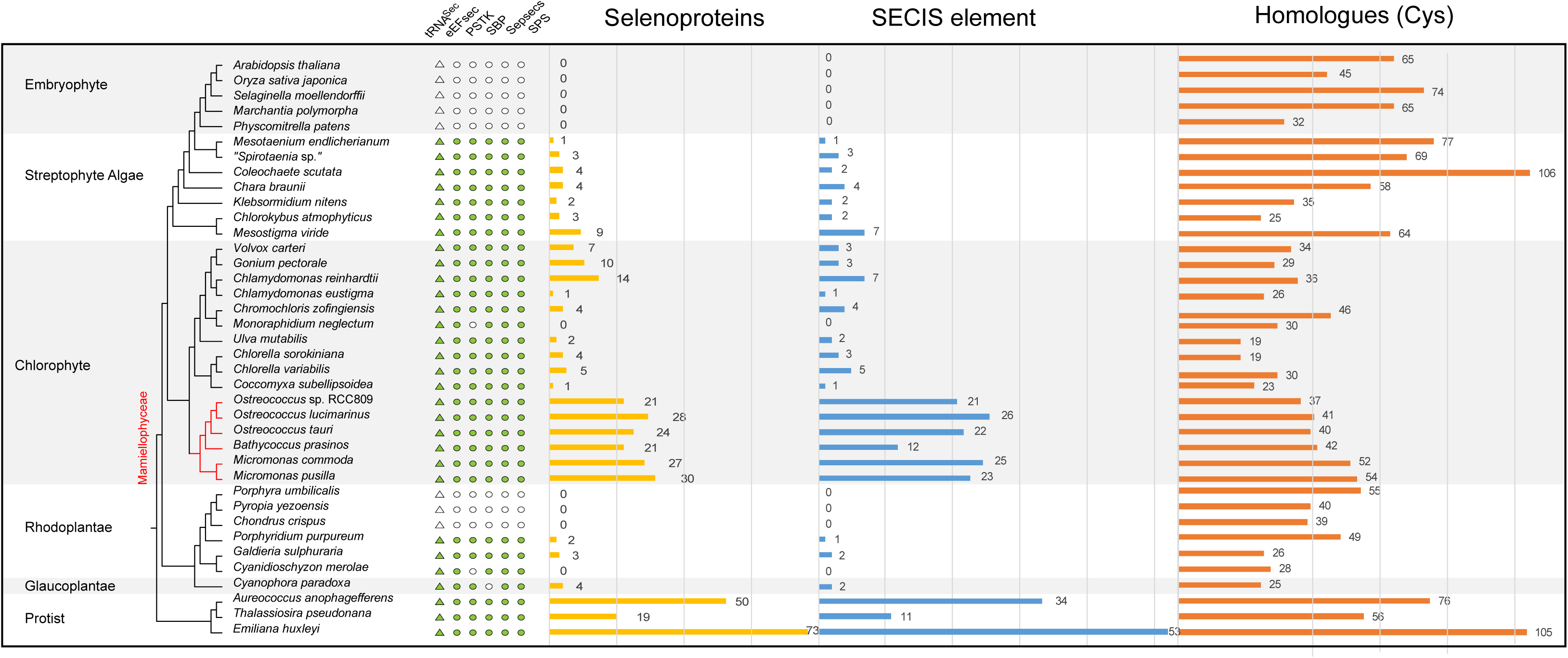
The number and distribution of selenoproteins, and enzymes involved in the Sec machinery. The phylogenetic tree was retrieved from the NCBI taxonomy database and the 1 KP Project (http://www.onekp.com). Presence (green symbols) or absence (empty symbols) of the enzymes involved in the Sec machinery (circles) and tRNASec (triangles) across sequenced embryophyte, streptophyte algae, chlorophyte, Rhodoplantae, Glaucoplantae and protist genomes are shown in the left panel. The distribution and number of selenoproteins are plotted in the yellow column in the second panel, and the predicted SECIS elements are represented by the blue bars. Distribution and number of selenoprotein homologues (Cys) are plotted in an orange column on the right panel. Prasinophyte algae (Mamiellophyceae) are highlighted in red.

**Figure 2.**
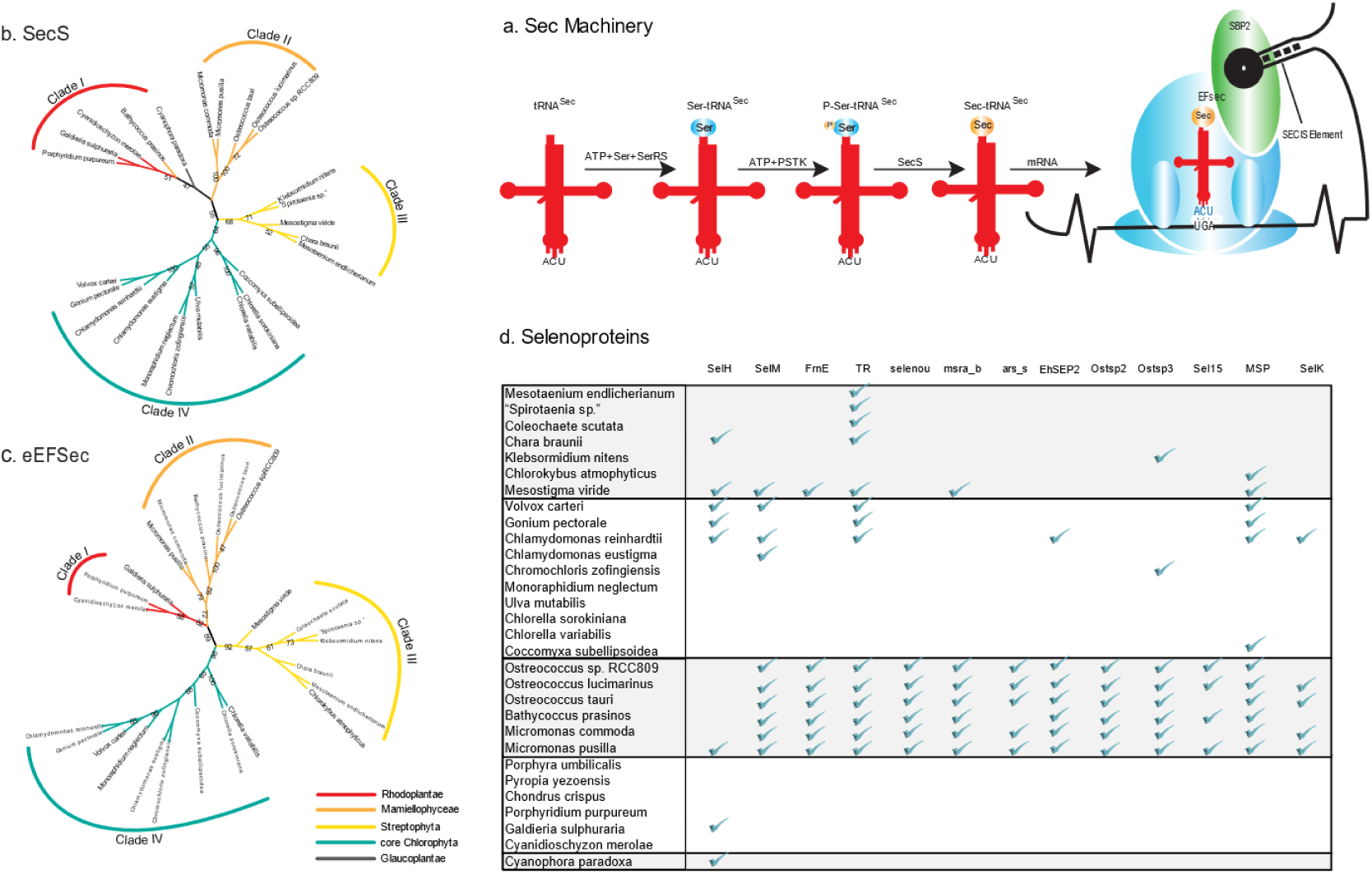
Phylogenetic analysis of enzymes involved in the Sec machinery. (a) Schematics of the selenoprotein biosynthesis pathway. (b-c) Maximum-likelihood trees of EFsec and SecS respectively. Bootstrap values >50% are shown. The tree support for internal branches was assessed using 500 bootstrap replicates. (d) Distribution of selected selenoproteins across the Archaeplastida. Presence of selenoproteins are shown by green check marks.

Although selenoproteins constitute only a small fraction of the proteome in any living organism, they play important roles in redox regulation, antioxidation, and thyroid hormone activation in animals including humans [16]. Sec incorporation has been well documented in animals, bacteria, and archaea, while the largest selenoproteome was reported in algae. In the pelagophyte alga *Aureococcus anophagefferens*, 59 selenoproteins were identified in its genome, compared with 25 selenoproteins in humans [17,18]. The green alga *Chlamydomonas reinhardtii* has at least ten selenoproteins, whereas the picoplanktonic, marine green alga *Ostreococcus lucimarinus* harbors 20 selenoprotein genes in its genome [3,19]. Considering that Se is essential for growth in at least 33 algal species that belong to six phyla, Sec incorporation is thought to be universal in diverse algal lineages [20]. In a previous study, no selenoproteins were found in any land plants [7], suggesting a complete loss of Sec incorporation after streptophyte terrestrialization. Exploring the Sec machinery across the Archaeplastida, especially in algae, would provide insight into its evolutionary dynamics in this important lineage of photosynthetic eukaryotes. Here in this study, we searched 38 plant genomes, including 33 algal species that represent the major algal lineages, for the Sec machinery and selenoproteins.

## 2. Results

### 2.1 Sec machinery in algae

To cover the plant tree of life, we selected 33 genomes of algal species and 5 embryophyte species with a focus on Archaeplastida, the major group of photosynthetic eukaryotes with primary plastids. The 33 algal species include one glaucophyte, six rhodophytes, 16 chlorophytes, and seven streptophyte algae. Another three species, the pelagophyte *A. anophagefferens*, the diatom *Thalassiosira pseudonana*, and the coccolithophorid *Emiliania huxleyi*, were also included to represent other distinct algal lineages (Supplementary Table S1A).

The Sec machinery was searched in 38 genome assemblies using Selenoprofiles [21] (See Methods). As shown in Figure 1, embryophytes lack the entire Sec machinery as previously reported [7]. Interestingly, the Sec machinery is not intact in all tested algal species. Among 33 algae, three Rhodoplantae lack the entire Sec machinery as in embryophytes. The chlorophyte *Monoraphidium neglectum*, and the rhodophyte *Cyanidioschyzon merolae* lack PSTK, and the glaucophyte *Cyanophora paradoxa* SBP2. According to the species tree, it seems that the Sec machinery was lost completely in one rhodophyte clade that includes *Porphyra umbilicalis, Pyropia yezoensis*, and *Chondrus crispus* and partially in a few other algal species (Figure 1).

### 2.2 Sec incorporation in the major algal lineages

In addition, we also identified the complete Sec machinery in some Rhodoplantae (Figure 1). The Rhodoplantae are often classified at the subphylum level into two clades, Cyanidiophytina and Rhodophytina [22], the latter consisting of 6 classes that can be grouped into two lineages: SCRP (Stylonematophyceae, Compsopogonophyceae, Rhodellophyceae, Porphyridiophyceae) and BF (Bangiophyceae, Florideophyceae) [23,24]. The entire Sec machinery was absent in *Porphyra, Pyropia* and *Chondrus* that belong to the BF clade (Figure 1).

The three Stramenopiles and haptophyte algal species encoded the complete Sec machinery, and generally also displayed more selenoproteins than most green algae [3,5,17,19,25]. In the Chlorophyta, the picoplanktonic Mamiellophyceae stand out because they not only encode the complete Sec machinery but also contain a large number of selenoproteins (Figure 1). In the remaining Chlorophyta comprising the three classes Trebouxiophyceae, Ulvophyceae and Chlorophyceae (the TUC clade according to [26]), except for *M. neglectum*, all other sequenced genomes encode the full Sec machinery and contain selenoproteins, although their number is considerably lower than in the Mamiellophyceae (Figure 1) supporting a previous report [5]. The number of selenoproteins among Chlorophyta is variable; very low numbers were encountered in *Chlamydomonas eustigma* and *Coccomyxa subellipsoidea*, the first isolated from acid mine drainage with very high sulfate content (and in this aspect resembling the cyanidiophyte *Galdieria sulphuraria* which also only has a few selenoproteins, Figure 1), the latter exclusively occurring in subaerial habitats (damp rocks and stones, [27]).

### 2.3 Variable number of selenoproteins identified in algae

Selenoproteins were scanned in the 38 plant genome assemblies using Selenoprofiles (Supplementary Figure S2), and their SECIS elements were identified in the 6-kb downstream of their putative stop codons by SECISearch3 [21,28]. There are some predicted selenoproteins that did not predict SECIS elements in the downstream region, especially in Mamiellophyceae, e.g. *Bathycoccus prasinos*, which may be because of lineage-specific characteristics or incomplete assembly [28]. The presence of selenoproteins in each assembled genome agrees with the intactness of the Sec machinery. In the rhodophyte clade that lacks the machinery or in the algae that miss one of the components, none of the known selenoproteins and SECIS elements were found in their genomes (except for *C. paradoxa*, in which the unidentified SBP2 protein may be incompletely assembled or other proteins replace the function of SBP2).

The Sec machinery is absent in embryophytes including the liverwort *Marchantia polymorpha* and the moss *Physcomitrella patens* (Figure 1). The availability of genomes (or transcriptomes) of all major lineages of streptophyte algae, the phylogeny of which can now be regarded as basically resolved [29], allowed identification of the likely step in the evolution of streptophytes when the loss of the Sec machinery and of selenoproteins occurred. As a first attempt to address this question, we searched the transcriptomic data from the 1KP project (http://www.onekp.com) for the presence of the Sec machinery and selenoproteins (268 algal species, 70 species of non-vascular (liverworts, mosses, hornworts) plants, and 175 species of monilophytes, lycophytes, and conifers). The number of enzymes of the Sec machinery and the number of selenoproteins were computed for each group (Supplementary Table S2). The sec machinery was completely absent from hornworts with no Sec incorporation machinery enzyme and selenoproteins. In liverworts and mosses, only a few selenoproteins were detected (2 and 1 respectively), and only a few enzymes of the Sec machinery were randomly distributed (in no bryophyte species were more than two of the five components of the Sec machinery detected: PSTK and SecS were absent in hornworts and eEFsec and SPS were absent in mosses) (Supplementary Table S2). In vascular plants, the Sec machinery was absent in all transcriptomes of all plants and no selenoproteins were detected (Supplementary Table S2). In the sister group of embryophytes, the Zygnematophyceae, enzymes of the Sec machinery were more widely distributed compared to bryophytes (Supplementary Table S2). In Zygnematophyceae, none among the five genes of the Sec machinery was found in their transcriptomes (four of the five components of the Sec machinery were present in about one third of the 40 taxa). It might be a consequence of the fragmentary nature of transcriptomes (e.g. we could not detect a complete Sec machinery in the transcriptomes of “*Spirotaenia* sp.” and *Mesotaenium endlicherianum*, although in both genomes the complete Sec machinery had been identified, Figure 1). Furthermore, selenoproteins were identified in only 15 of the 40 Zygnematophyceae and their number per species was low. Again, we did not detect selenoproteins in the transcriptomes of “*Spirotaenia sp.*” and *M. endlicherianum*, although in their genomes a few genes encoding selenoproteins were identified (note that the number of selenoproteins, as well as components of the Sec machinery, is higher in “*Spirotaenia* sp.” because of its recent genome triplication; Cheng et al. unpublished observ). In the other clades of the streptophyte algae (Coleochaetophyceae, Charophyceae, Klebsormidiophyceae and Mesostigmatophyceae) the situation is similar to that in Zygnematophyceae, a complete Sec machinery is present but the number of selenoproteins identified is low, especially in the subaerial taxa (two in *Klebsormidium nitens* and three in *Chlorokybus atmophyticus*), the only exception being the scaly flagellate *Mesostigma viride* with 9 identified selenoproteins (Table 1 and Supplementary Table S2).

**Table 1.**
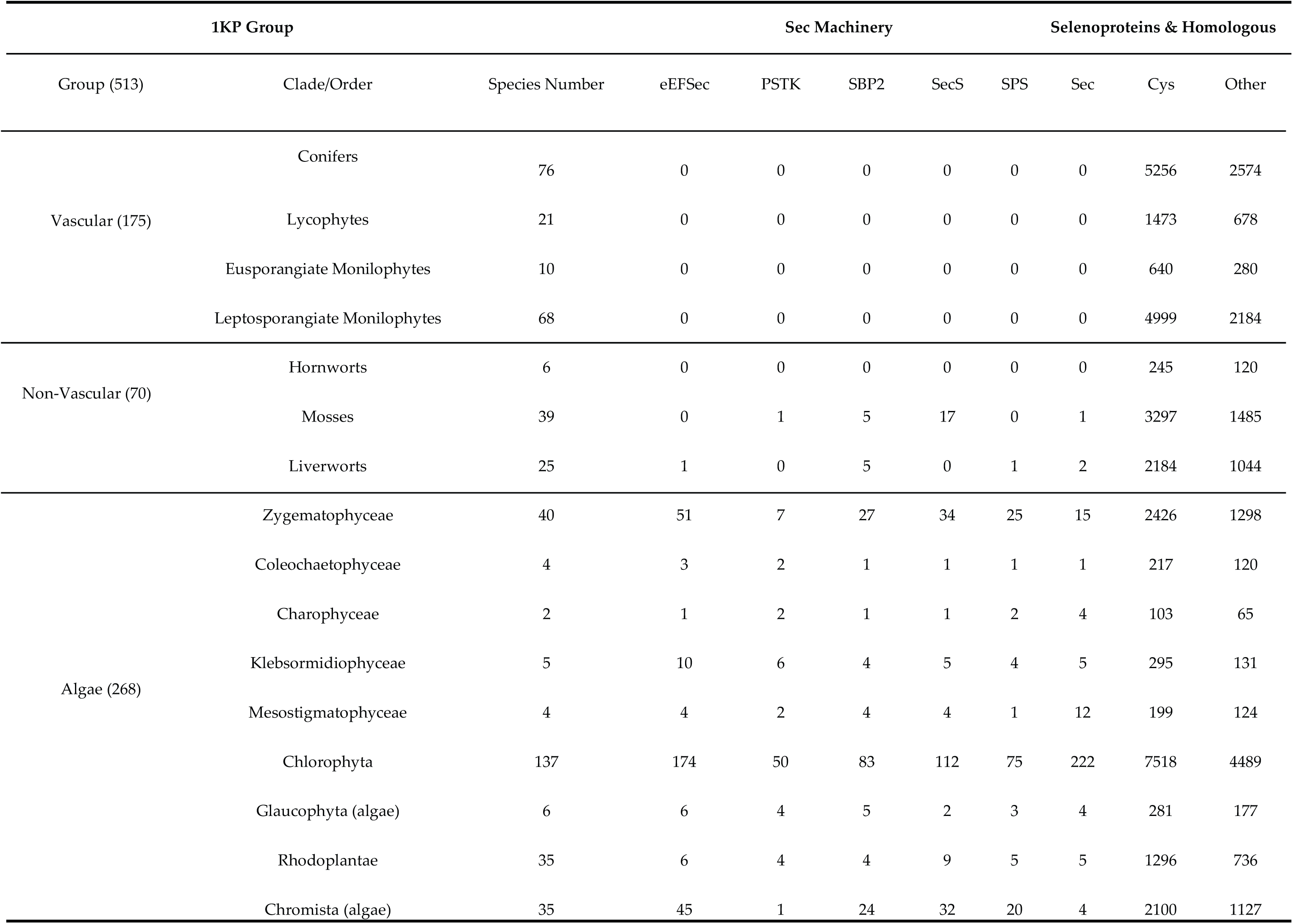
Number of enzymes involved in the Sec incorporation machinery and selenoproteins. The number of enzymes of the Sec incorporation machinery and selenoproteins are detected by Selenoprofiles across the sequenced Algae, Liverworts, Mosses, Hornworts and a part of lower embryophyte genomes and transcriptomes (from the 1 KP project).

### 2.3 Phylogenetic analysis of the enzymes involved in the Sec machinery

To further analyze the evolution of the Sec machinery, we conducted phylogenetic analyses of five genes encoding Sec-containing enzymes from the available Archaeplastida genome data set. The phylogenetic trees of PSTK, SBP2, and SPS showed either insufficient phylogenetic signal resulting in low support values for internal branches (PSTK) or very long branches in several taxa (SBP2, SPS) that led to spurious topologies due to long-branch attraction or indicated discordant gene histories (Supplementary Figure S3 a, b and c).

The phylogenies of EFsec and SecS were largely congruent with some support for internal branches (especially EFsec) that roughly corresponded to the known phylogenetic relationships among higher order taxa, although relationships within some groups (e.g. streptophyte algae) remained unresolved (Figure 2b and 2c). The EFsec phylogeny revealed four clades of sequences that were reasonably well supported: clade I comprised three sequences of Rhodoplantae, clade II six sequences of picoplanktonic Mamiellophyceae, clade III seven sequences of streptophyte algae, and clade IV 9 sequences from the TUC clade (3 sequences of Trebouxiophyceae and 6 sequences of Chlorophyceae).

#### 2.3.1 Phylogenetic analysis of eukaryotic SPS proteins

We built an SPS gene set comprising both prokaryotes and eukaryotes to reconstruct a global SPS phylogenetic tree (Figure 3). SPS split into three well-separated clades: Clade I including a diverse range of bacteria, most of the Viridiplantae, and protists with secondary plastids (Stramenopiles, cryptotphytes, haptophytes and Apicomplexa), clade II containing bacteria and four species of green algae (*Chara braunii*; *Gonium pectorale*; *C. reinhardtii*; and *Volvox carteri*), and Clade III including archaea, a diverse range of protists (photosynthetic and non-photosynthetic), fungi, and three rhodophytes but no other Archaeplastida (Supplementary Table S3). The sequence of SPS Clade I contains three domains: Pyr_redox_2, AIRS and AIRS_C. However, sequences of Clade II and Clade III only showed the presence of AIRS and AIRS_C. The SPSs from Clade II and Clade III have different characteristics of domain arrangements (Supplementary Figure S3; as the phylogenetic tree suggested, potential horizontal gene transfer might have occurred in Clades I and II.). The SPS of the three Volvocales (*C. reinhardtii, G. pectorale* and *V. carteri*) from Clade II might have been acquired by horizontal gene transfer (HGT) from cyanobacteria, because they form a monophylum (92% boostrap support) with two terrestrial, filamentous cyanobacteria (*Tolypothrix bouteillei, Scytonema hofmannii*) which are themselves nested within a larger radiation of bacteria (Supplementary Figure S3). For *C. braunii*, we suspect that this gene derived from either a (cyano) bacterial or volvocalean contamination.

**Figure 3.**
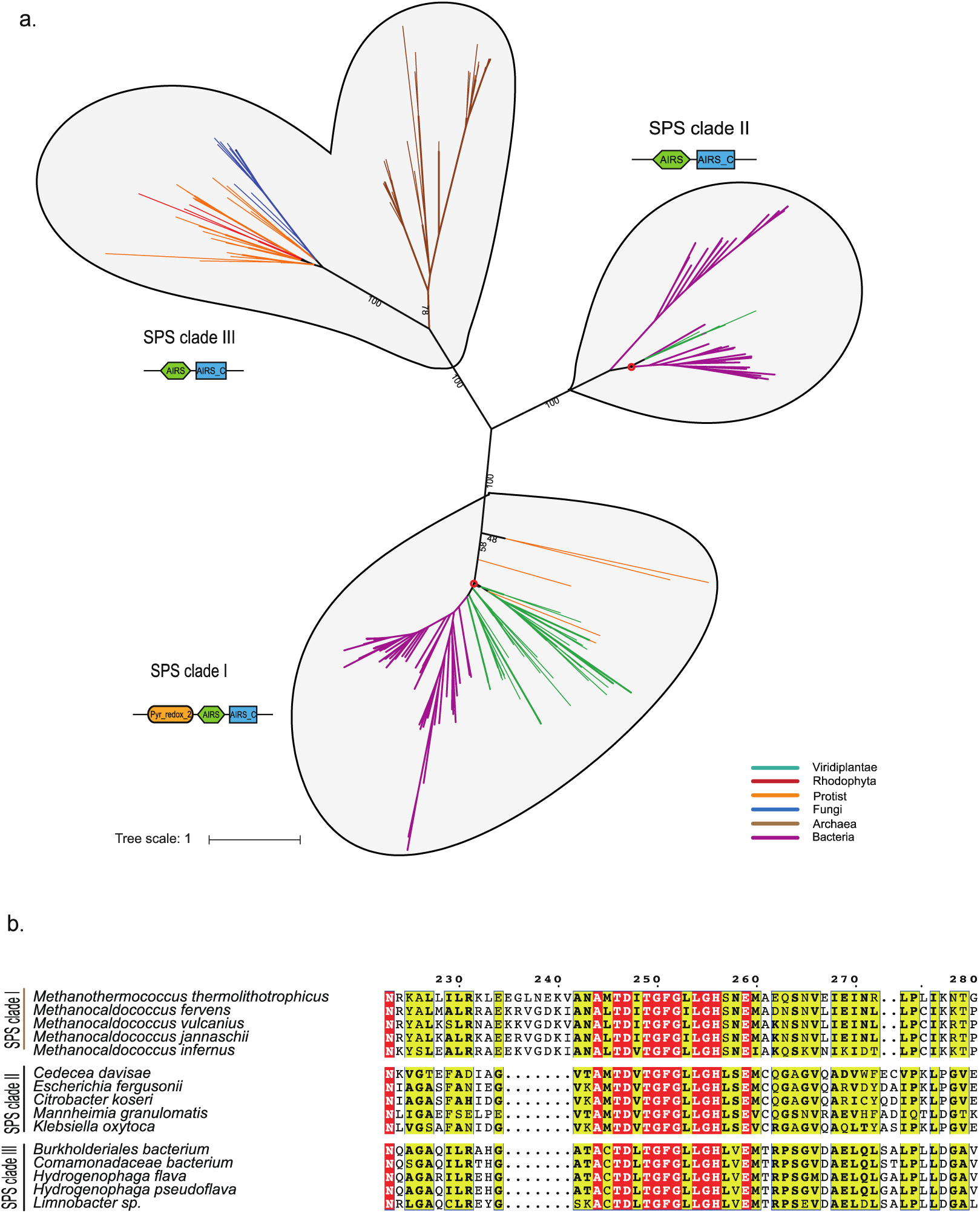
Phylogenetic analysis of selenophosphate synthetase (SPS). (a) Reconstructed protein phylogeny of the reference set of SPS proteins. The red point denotes potential horizontal gene transfer events in SPS clades I and II. (b) Alignment of SPS domains of the three SPS clades.

### 2.4. Distribution of types of selenoproteins among Archaeplastida

A comprehensive analysis of the distribution of selenoproteins revealed that picoplanktonic Mamiellophyceae possess an expanded set of selenoproteins, whereas some selenoproteins had a scattered distribution among other Archaeplastida (Figure 2d, and Supplementary Figure S2). This may be related to the distinct types of eEFsec and SecS present in the Mamiellophyceae (Figure 2b, and Figure 2c). Functional annotation of the selenoproteins in the genomes of the Mamiellophyceae showed that they are mainly involved in oxidative stress response and adaptation. The MsrA selenoprotein, e.g., is a key Sec-containing enzyme for the repair of oxidatively damaged peptides. However, MsrA_b, a bacterium-like MsrA selenoprotein, was identified only in the picoplanktonic Mamiellophyceae and in *M. viride* (Figure 2d), suggesting that early-diverging lineages of aquatic Viridiplantae might be subjected to stronger oxidative stress, and MsrA_b but not MsrA (Supplementary Figure S2) is essential for these species to perform the repair of peptides. Another Sec-containing oxidoreductase (FrnE) is present in the Mamiellophyceae and in *M. viride* but not in any other Archaeplastida genome sequenced (Figure 2d). FrnE is a cadmium-inducible protein that is characterized as a disulfide isomerase having a role in oxidative stress tolerance. Therefore, it also supports the above hypothesis that Mamiellophyceae and *M. viride* (or perhaps scaly green algae, in general) need these enzymes to cope with stronger oxidative stress. In this context, it is interesting to note that in the bloom-forming pelagophyte alga *A. anophagerfferens*, which has the second largest number of selenoproteins reported (50), a large number of redox active selenoproteins were overexpressed upon infection by a giant virus of the Mimiviridae clade [30], which suggests that viral infections, that are also prominent in the picoplanktonic Mamiellophyceae (prasinoviruses; [31]) and have also been described in *M. viride* [32], may elicit similar responses in their hosts. Viral infections are unknown in the three Volvocales studied (*C. reinhardtii, V. carteri*, and *G. pectorale*), however Volvocales are often subject to invasion by parasitic protists or fungi [33-35] and this could perhaps explain the presence of selenoproteins in these taxa.

## 3. Discussion

### 3.1 The distribution of the Sec machinery and selenoproteins in algae

It has been hypothesized that the Sec machinery and selenoproteins were lost in Viridiplantae upon transfer from an aquatic to a terrestrial environment perhaps related to the paucity of a suitable chemical species of selenium (i.e. selenite) in most terrestrial environments [7,9,20,36-38,40]. The results presented here support this notion and further suggest that the Sec machinery was lost in the common ancestor of embryophytes as all extant embryophytes lack this machinery in their genomes (Figure 1). The few enzymes of this machinery that were detected in the transcriptomes of some liverworts and mosses (Supplementary Table S2) likely represent contaminations. Interestingly, although the complete Sec machinery is still present in all classes of streptophyte algae, the number of selenoproteins detected in the subaerial species (*C. atmophyticus, K. nitens, “Spirotaenia* sp.*”, M. endlicherianum*) was low (1-3 proteins), whereas in the aquatic species (*M. viride, C. braunii, C. scutata*) more selenoproteins (4-9 proteins) were found (Figure 1). Very low numbers of selenoproteins (i.e. one protein) were also encountered in subaerial/acidophilic species of Chlorophyceae (*C. eustigma, C. subellipsoidea*) and in the subaerial/acidophilic Rhodoplantae (*G. sulphuraria*). These results corroborate the hypothesis that adaptation to subaerial/terrestrial or acidophilic habitats supports the gradual loss of selenoproteins in diverse groups of algae. We suspect that once selenoproteins have been lost, selection on maintaining the Sec machinery is abolished. Intermediate stages in this process may be seen in the subaerial chlorophyte *M. neglectum* (now *M. braunii*) and in the acidophilic red alga *C. merolae* [36], which each lost one enzyme (PSTK or SBP respectively) of the Sec machinery. We hypothesize that once the Sec machinery is lost, transfer of algae to aquatic (marine) habitats (as in most species of Rhodoplantae) will not lead to reappearance of selenoproteins (some red algae exposed to strong oxidative stress such as *P. umbilicalis* have developed intimate associations with bacteria that express selenoproteins [40]). Similarly, transcriptomes of later-diverging Zygnematophyceae (i.e. Desmidiales), that are predominantly aquatic in mostly acidic environments (bogs), also either lack selenoproteins or have only 1-2 selenoprotein(s) (Supplementary Table S2). It will be interesting to learn, once their genome sequences will become available, whether they display a Sec machinery or not. Palenik et al. [19] proposed a trade-off between increased Se requirements but decreased nitrogen requirements for peptide synthesis in *Ostreococcus spp.*, and it is worth noting that this genus encodes a surprisingly high number of selenocysteine-containing proteins relative to its genome size [19]. The core Chlorophyta showed a similar number of genes involved in nitrogen metabolism as the picoplanktonic Mamiellophyceae (Supplementary Table S4). In Trebouxiophyceae and Ulvophyceae (represented by *Ulva mutabilis*), fewer selenoproteins were identified than in the Mamiellophyceae. Functional annotation of the selenoproteins in Trebouxiophyceae and Ulvophyceae showed that they mainly participated in some redox activities such as redox signaling (thioredoxin reductase, TR) and oxidative stress response (glutathione peroxidase, GPx) (Supplementary Figure S2). However, it is still unclear why Trebouxiophyceae and Ulvophyceae possess fewer selenoproteins, the first occur in freshwater or are often subaerial, the latter is mostly multicellular and may not require the diversity of highly reactive selenoenzymes characteristic for picoeukaryotes.

### 3.2 Probable Horizontal Gene Transfer of SPS and some selenoproteins

SPS was detected in both prokaryotes and eukaryotes, although their sequence similarity is quite low (∼30%; [4]). Our phylogenetic analyses resolved three clades of SPS genes with mixed species composition of prokaryotes and eukaryotes suggesting HGT among these unrelated organisms. For SPS Clade II, we provided evidence that a single HGT event occurred from terrestrial cyanobacteria into the common ancestor of *C. reinhardtii, V. carteri*, and *G. pectorale*. Several selenoproteins of the picoplanktonic Mamiellophyceae may also have had their origin in the domain bacteria and been recruited from bacteria (perhaps via viruses) through HGT. Selenoproteins are relatively common in bacteria, about 34% of the sequenced bacteria utilize Sec, mostly different groups of proteobacteria [38, 39]. Phylogenetic analyses of selenoproteomes in bacteria have identified rampant losses of selenoproteins but also occasional HGT events, even between domains (bacteria and archaea) [42,43]. It is tempting to speculate that these HGTs supported bloom-forming, marine microalgae that often lack cell walls, their cells being covered only by mineralized or non-mineralized scales, to cope with viral invasions using their highly redox-reactive selenoproteins.

## 4. Conclusion

A phylogenomic analysis of the selenocysteine (Sec) –machinery and selenoproteins in genomes and transcriptomes of diverse Archaeplastida provided evidence for complete or partial loss of the Sec-machinery in several, unrelated lineages accompanied by loss of selenoproteins. In streptophytes, the Sec-machinery and selenoproteins were apparently lost in the common ancestor of embryophytes, as the Sec-machinery was present in all lineages of streptophyte algae but absent in embryophytes. The number of selenoproteins identified in algae correlated with the type of their habitats, low numbers of selenoproteins were encountered in algae thriving in subaerial/terrestrial or acidic environments. The large number of selenoproteins found in some bloom-forming, marine microalgae may be related to their function in the defense against viral infections. Some components of the Sec-machinery and selenoproteins may have been acquired by algae through horizontal gene transfer from bacteria.

## 5. Materials and Methods

### 5.1 Data information

A total of 38 genome sequences were used in this study, the genomes including 5 embryophytes, 7 streptophyte algae, 16 chlorophytes, 6 Rhodoplantae, 1 Glaucoplant and 3 photosynthetic protists (two stramenopiles and a haptophyte). The transcriptomes contained 121 green algae, 25 liverworts, 6 hornworts, 38 mosses, and 170 terrestrial plants (Supplementary Table S1A). The 33 whole genome assemblies were downloaded from the NCBI genome database. In addition, 5 newly assembled streptophyte algal genomes were used, including *Mesotaenium endlicherianum* (strain CCAC 1140), “*Spirotaenia* sp.” (strain CCAC 0220), *Coleochaete scutata* (strain SAG 110.80), *Mesostigma viride* (strain CCAC 1140), *Chlorokybus atmophyticus* (strain CCAC 0220). The CCAC strains were obtained from the Culture Collection of Algae at the University of Cologne (http://www.ccac.uni-koeln.de/). All cultures were axenic, and during all steps of culture scale-up until nucleic acid extraction, axenicity was monitored by sterility tests as well as light microscopy. Total RNA was extracted from *M. viride* using the Tri Reagent Method, and from *C. atmophyticus* using the CTAB-PVP Method as described in Johnson et al [44]. Total DNA was extracted using a modified CTAB protocol [45,46]. The phylogenetic backbone of algae was retrieved from the NCBI taxonomy database (https://www.ncbi.nlm.nih.gov/Taxonomy/CommonTree/wwwcmt.cgi). The completeness of genome assemblies was assessed by BUSCO 3.0.2 with eukaryote gene database [47]. The results were listed in the Supplementary Table S1B. We also counted the usage of stop codons for the single-copy genes. The results were shown in Supplementary Figure S1 (Supplementary Table S1B).

### 5.2 Sec incorporation machinery

The genome sequences were searched for the Sec incorporation machinery by the Selenoprofiles pipeline (version 3.0, http://big.crg.cat/services/selenoprofiles) with the parameter “-p machinery” [21,48]. Firstly, we ran the pipeline with profile-based Sec machinery. To reduce the incomplete gene sequence mistakes, the blastp version 2.6.0+ (e-value <10^-5^) was used against the predicted genes as in a special algae database to detect Sec machinery. In addition, transcriptome data were also searched using the same methods. First, the nucleic acid sequences were searched by Selenoprofiles, and then subjected to blastp (e-value <10^-5^) with the predicted algae-specific Sec machinery database.

### 5.3 Identification of the selenocysteine tRNA (tRNA^Sec^)

Secmarker version 0.4 (http://secmarker.crg.es/index.html) was used to identify the dedicated tRNA^Sec^ in the genome sequences [49]. The predicted secondary structure was drawn with the parameter “-plot”.

### 5.4 Prediction of selenoproteins and SECIS elements

Selenoproteins were identified from the genome assemblies with Selenoprofiles with the parameter “-p metazoa, protist, prokarya”. The candidates were filtered with cutoff: e-value <0.01 and the sensible AWSIc Z-score > −3. SECIS elements were searched in the 6-kb DNA sequences downstream of predicted selenoprotein genes at the SECISearch3 website (http://seblastian.crg.es/; with the parameter “-output_three_prime, -output_secis”) [28].

### 5.5 Phylogenetic tree construction

In phylogenetic analysis, each candidate was searched by Selenoprofiles and blastp version 2.6.0+ [44] to detect more candidates (e-value < 1 × 10-5). Multiple sequence alignments were performed by MAFFT version 7.310 [50,51]. In eEFSec, SecS, PSTK, and SBP2, the maximum-likelihood tree was constructed for each protein family using the IQ-TREE software with 500 bootstrap replicates [52]. The SPS maximum-likelihood trees were constructed for each protein family using the RAxML version 8.2.4 with the GTR+I+G model [53]. For the phylogeny of SPS (SelD), the bacteria sequences were downloaded from the NR database by submitting every alga SPS sequences to nr databases. All target bacterial sequences were retrieved but only several randomly chosen sequences in each bacterial phylum were used for the SPS phylogenetic analyses. Representative archaea and protist sequences were used in the analysis of SPS. In addition to this, the lately reported 9 fungi that utilize Sec were also added (192 sequences) [4,50].

### 5.6 Identification of conserved motifs and domains

Pfam 32.0 (http://pfam.xfam.org/) was used to identify the domains in the Sec incorporation machinery [54]. Additional motifs were identified by Multiple Em for Motif Elicitation 5.0.5 (MEME, http://meme-suite.org/). The alignment of the SPS domain was visualized by ESPript 3.0.

## Supporting information

http://db.cngb.org/cnsa

## Supplementary Materials

The sequences of selenoprotein which we identified from the green algae (*Mesostigma viride, Chlorokybus atmophyticus, Klebsormidium nitens, Chara braunii, Coleochaete scutata, “Spirotaenia* sp.*”, Mesotaenium endlicherianum*) are available in the CNGB Nucleotide Sequence Archive (CNSA: http://db.cngb.org/cnsa; accession number CNP0000452). The specific details regarding other genes which were used in this study are available in supplementary File S5.

## Author Contributions

Data curation, Yan Xu and Linzhou Li; Formal analysis, Hongping Liang, Hongli Wang and Haoyuan Li; Funding acquisition, Xin Liu and Huan Liu; Investigation, Hongping Liang; Methodology, Linzhou Li, Gengyun Zhang and Sibo Wang; Project administration, Tong Wei and Huan Liu; Resources, Michael Melkonian; Software, Yan Xu and Linzhou Li; Supervision, Sunil Kumar Sahu, Xin Liu, Sibo Wang and Huan Liu; Visualization, Hongping Liang; Writing – original draft, Sunil Kumar Sahu and Sibo Wang; Writing – review & editing, Sunil Kumar Sahu, Xian Fu and Michael Melkonian.

## Acknowledgments

We thank Dr. Shifeng Cheng for kindly providing the gene sequences of “*Spirotaenia sp.*” and *Mesotaenium endlicherianum*.

## Abbreviations

Sec: Selenocysteine
Se: Selenium
SECIS: Selenocysteine Insertion Sequence
PSTK: O-phosphoseryl-transfer tRNA^Sec^ kinase
SecS: Sec synthase
SPS: selenophosphate synthetase 2
SBP2: SECIS-binding protein 2
eEFSec: Sec-specific elongation factor
CTAB: Cetyl trimethylammonium bromide

## Supplementary Figures

**Supplementary Figure S1: Completeness of genome assemblies and Stop codon statistics.** The genome quality was assessed by the BUSCO program. The number of single, multiple, missing and fragmented genes are shown in the middle histogram. The stop codon usage of genes is shown on the right histograms. The species that lost the Sec machinery are marked with an asterisk in the tree. Each group was colored in different background in the left column.

**Supplementary Figure S2: Species tree with families of selenoproteins and homologues (Cys) by Selenoprofiles.** Each column corresponds to a selenoprotein family plotted in different color to distinguish Sec-containing (green), Cys homologues (red), and others (grey). 33 out of 62 selenoproteins families having at least one Sec-containing gene were selected for the analysis. The number in the left box represents the number of selenoproteins.

**Supplementary Figure S3: The phylogenetic tree of PSTK, SBP2, and SPS.** All the predicted candidates were used to reconstruct a maximum-likelihood tree, with 500 bootstrap replicates. Here, Orange: Mamiellophyceae; Yellow: core Chlorophyta; Green: Streptophyte algae; Red: Rhodoplantae; and Grey: Glaucoplantae.

**Supplementary Figure S4: The phylogenetic tree of SPS.** The maximum-likelihood tree by using the reference set of SPS proteins was reconstructed using the GTR+I+G model.

**Supplementary Figure S5: The alignment of the SPS domain**. The complement alignment of SPS domains in the three clades. Nucleotide conservation is indicated with red and yellow coloring. The three clades are separated by spaces. From top to bottom, they are clade III, clade II and clade I.

## Supplementary Table

**Supplementary Table S1:** Table S1A: The contents of each column are species class, names, assembly identifiers and BUSCO assesses genome quality of genomes analysed in this study. Table S1B: The distribution of stop codon in each species.

**Supplementary Table S2:** The number of enzymes of the Sec machinery and the number of selenoproteins were computed for each group.

**Supplementary Table S3:** The details of species in SPS phylogenetic tree: SuperKingdom, Phylum/Class, GI Number, Species Name and SPS Clade.

Supplementary Table S4: Genes important for Nitrogen metabolism pathway in Chlorophyta.

