## Supplementary material for "Phylogenomics provides new insights into gains and losses of selenoproteins among Archaeplastida": http://db.cngb.org/cnsa

**Supplementary Figure S1: Completeness of genome assemblies and Stop codon statistics.** The genome quality was assessed by the BUSCO program. The number of single, multiple, missing and fragmented genes are shown in the middle histogram. The stop codon usage of genes is shown on the right histograms. The species that lost the Sec machinery are marked with an asterisk in the tree. Each group was colored in different background in the left column.

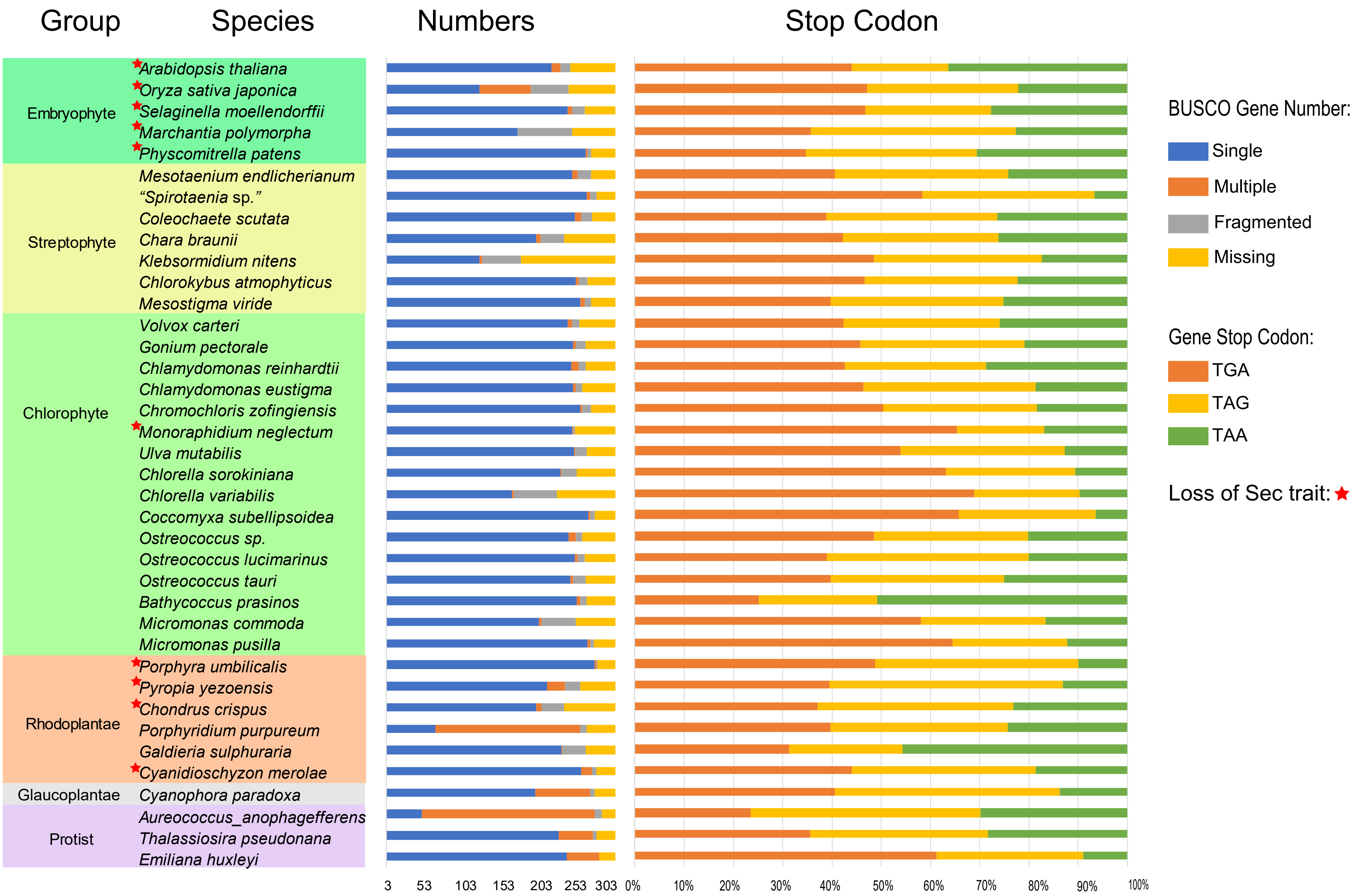

**Supplementary Figure S2: Species tree with families of selenoproteins and homologues (Cys) by Selenoprofiles.** Each column corresponds to a selenoprotein family plotted in different color to distinguish Sec-containing (green), Cys homologues (red), and others (grey). 33 out of 62 selenoproteins families having at least one Sec -containing gene were selected for the analysis. The number in the left box represents the number of selenoproteins.

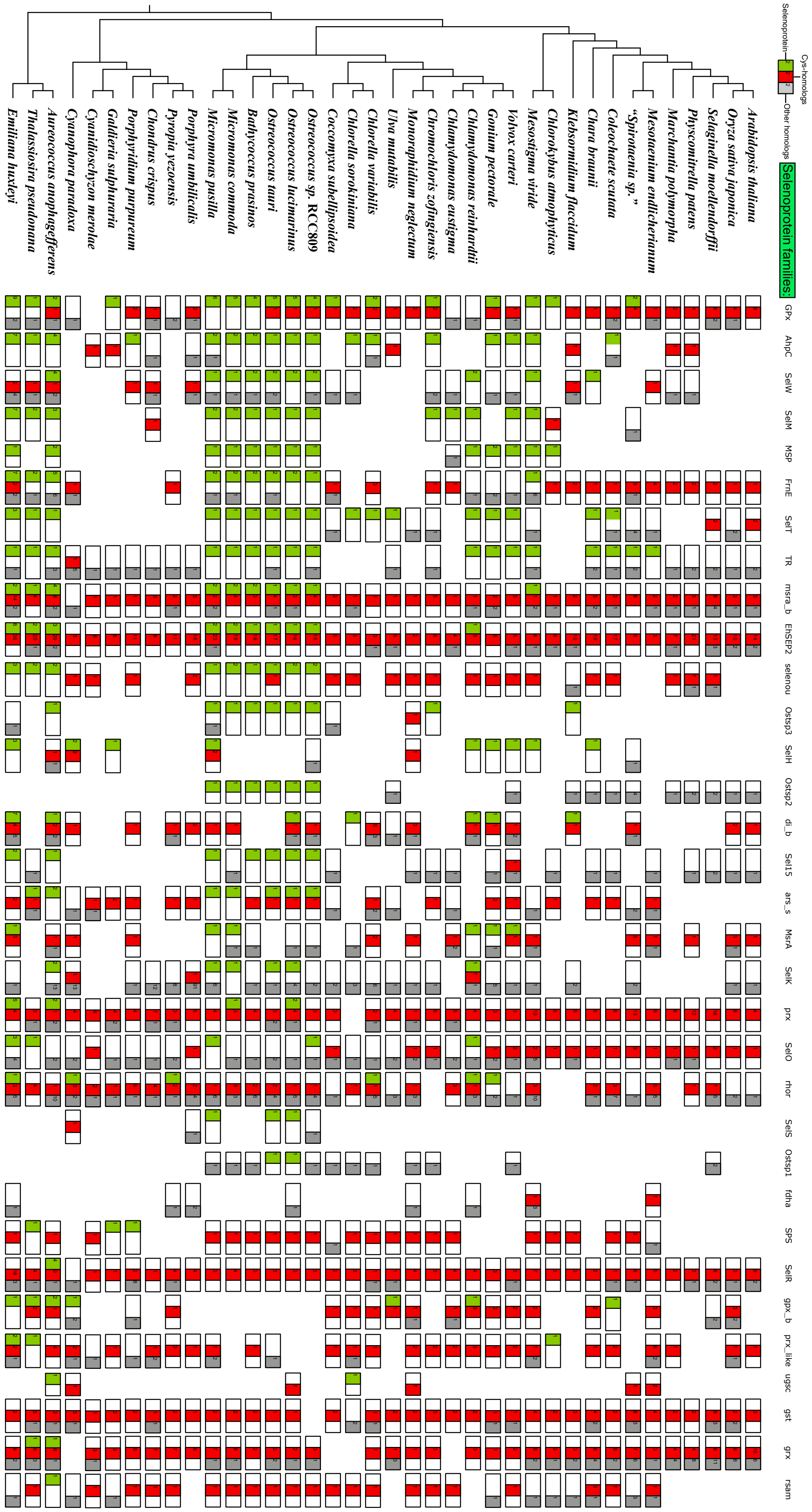

**Supplementary Figure S3: The phylogenetic tree of SBP2, PSTK and SPS.** All the predicted candidates were used to reconstruct a maximum-likelihood tree, with 500 bootstrap replicates. Here, Orange: Mamiellophyceae; Yellow: core Chlorophyta; Green: Streptophyte algae; Red: Rhodoplantae; and Grey: Glaucoplantae.

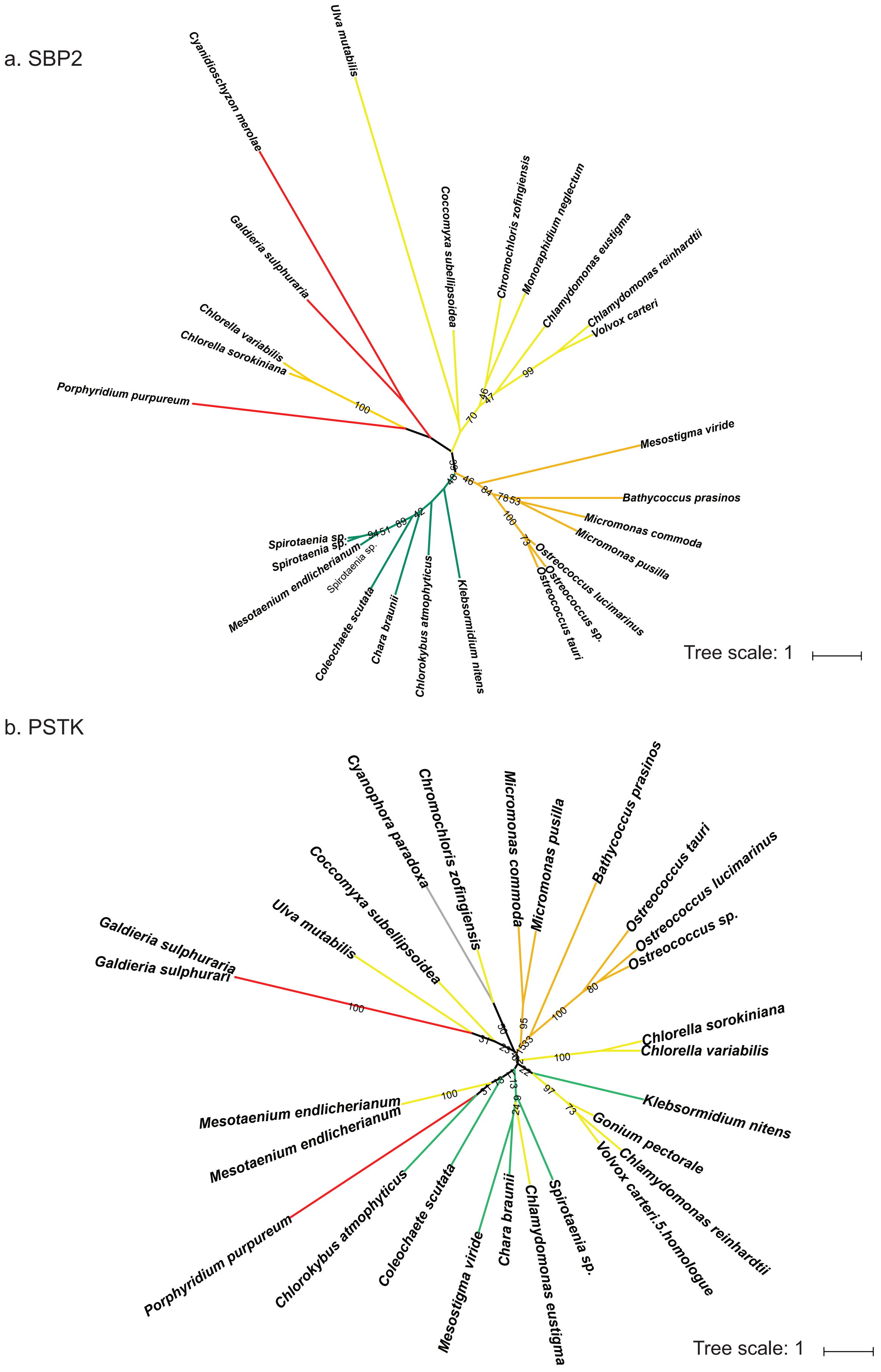

c. SPS

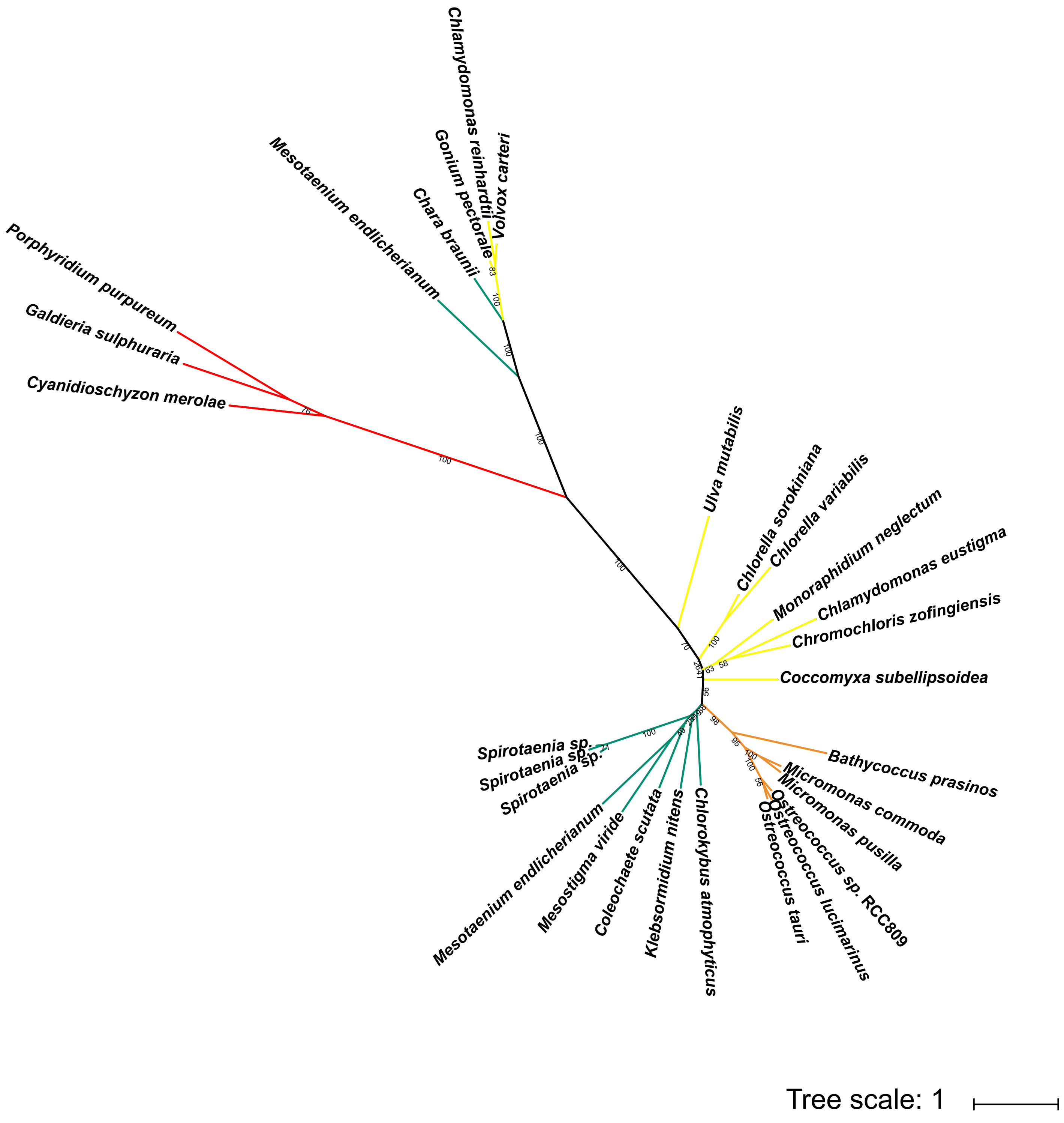
Chlamydomonas reinhardtii
Volvox carteri
Gonium pectorale
Chara braunii
Mesotaenium endlicherianum
Porphyridium purpureum
Galdieria sulphuraria
Cyanidioschyzon merolae
Ulva mutabilis
Chlorella sorokiniana
Chlorella variabilis
Monoraphidium neglectum
Chlamydomonas eustigma
Chromochloris zofingiensis
Coccomyxa subellipsoidea
Bathycoccus prasinos
Micromonas commoda
Micromonas pusilla
Ostreococcus sp. RCC809
Ostreococcus lucimarinus
Ostreococcus tauri
Chlorokybus atmophyticus
Klebsormidium nitens
Coleochaete scutata
Mesostigma viride
Mesotaenium endlicherianum
Spirotaenia sp.
Spirotaenia sp.
Spirotaenia sp.
Tree scale: 1

**Supplementary Figure S4: The phylogenetic tree of SPS.** The maximum-likelihood tree by using the reference set of SPS proteins was reconstructed using the GTR+I+G model.

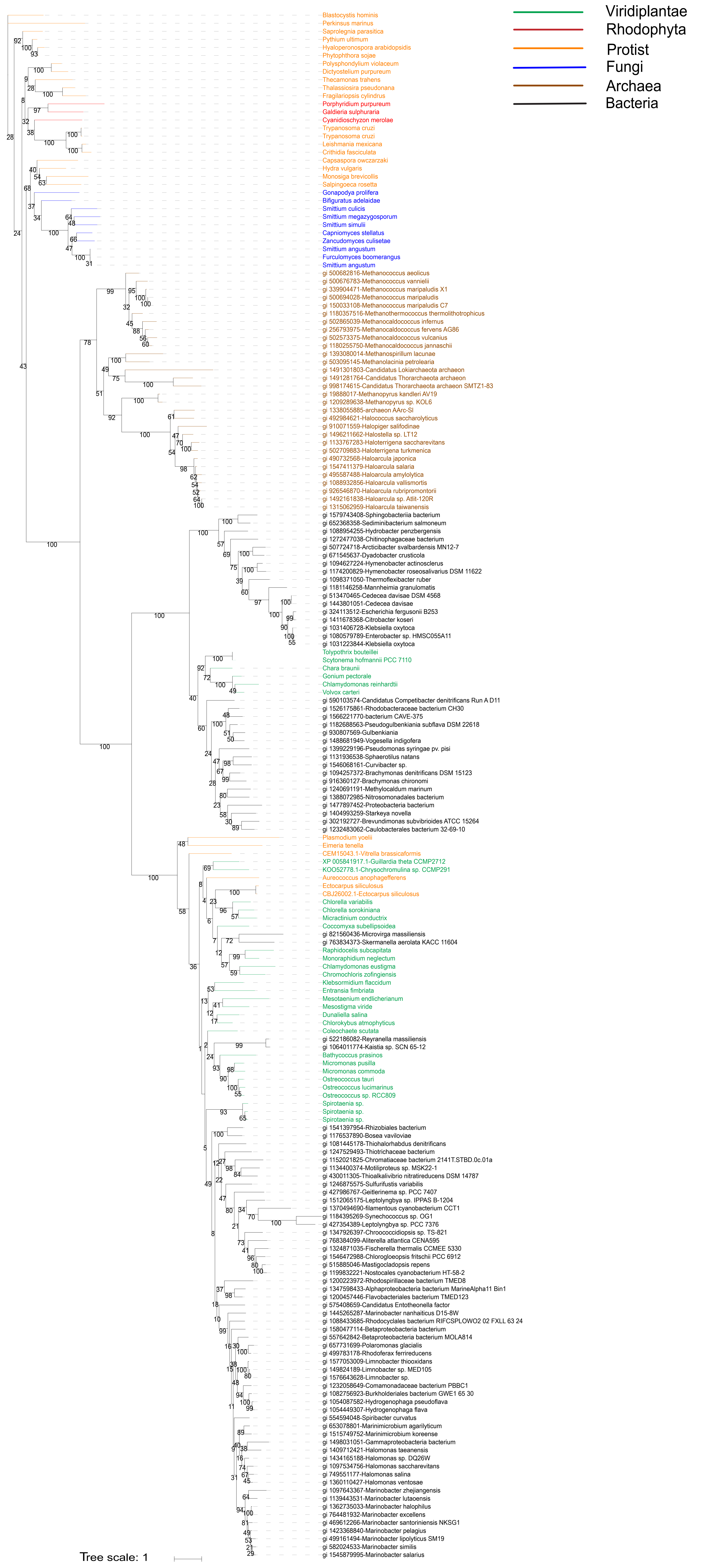

**Supplementary Figure S5: The alignment of the SPS domain.** The complement alignment of SPS domains in the three clades. Nucleotide conservation is indicated with red and yellow coloring. The three clades are separated by spaces. From top to bottom, they are clade III, clade II and clade I.

```
Methanothermococcus thermolithotrophicus  ............................................................
Methanocaldococcus fervens                ............................................................
Methanocaldococcus vulcanius              ............................................................
Methanocaldococcus jannaschii             ............................................................
Methanocaldococcus infernus               ............................................................

Cedecea davisae                           ............................................................
Escherichia fergusonii                    ............................................................
Citrobacter koseri                        ............................................................
Mannheimia granulomatis                   ............................................................
Klebsiella oxytoca                        ............................................................

Burkholderiales bacterium                 DLVLIGGGHSHVGVLRMFAMKPEPGVRITVICTDIDTPYSGMLPGYISGHYSFDEVHIDL
Comamonadaceae bacterium                  DLVLVGGGHSHVVALRMLAMQPEPGLRITLVCTDIDTPYSGMLPGYVSGHYSFDEVHIDL
Hydrogenophaga flava                      DLVLVGGGHSHVVALRMLAMRPEPGLRITMVCTDIDTPYSGMLPGYIAGHYSFDEVHIDL
Hydrogenophaga pseudoflava                DIVLVGGGHSHVGVLRMFGMKPWPGVRLTLICTDIHTPYSGMLPGYIAGHYSYDDVHIDL
Limnobacter sp.                           DLVLVGGGHSHVVVLRMLAMQPEPGLRITLICTDIDTPYSGMLPGYISGHYSFDDVHIDL


Methanothermococcus thermolithotrophicus  ............................................................
Methanocaldococcus fervens                ............................................................
Methanocaldococcus vulcanius              ............................................................
Methanocaldococcus jannaschii             ............................................................
Methanocaldococcus infernus               ............................................................

Cedecea davisae                           ............................................................
Escherichia fergusonii                    ............................................................
Citrobacter koseri                        ............................................................
Mannheimia granulomatis                   ............................................................
Klebsiella oxytoca                        ............................................................

Burkholderiales bacterium                 GRLCAYTGARFFHDAVVGIDRQNQKVICQNRPPVAYDLLSINIGSTPQVRHVAGAQSLAV
Comamonadaceae bacterium                  GRLAAFAGARFIHGEVTGLDRASQRVLLKGRPAIPYDLLSINTGSTPNVRQVDGAQAHAV
Hydrogenophaga flava                      GRLAVFAGARFIHGEVTGLDRANQRVLLKGRPSIPYDLLSINTGSTPNVRQVDGAQAHAV
Hydrogenophaga pseudoflava                GRLCAFAGARLFKDEVVGIDRANQKVICKNRPPVAYDALSINIGSTPQVQLVPGALENAV
Limnobacter sp.                           GRLAAFAGARFIHGEVTGLDRANQRVLLRDRPSVPYDLLSINTGSTPNVRQVDGARAHTV


Methanothermococcus thermolithotrophicus  ............................................................
Methanocaldococcus fervens                ............................................................
Methanocaldococcus vulcanius              ............................................................
Methanocaldococcus jannaschii             ............................................................
Methanocaldococcus infernus               ............................................................

Cedecea davisae                           ............................................................
Escherichia fergusonii                    ............................................................
Citrobacter koseri                        ............................................................
Mannheimia granulomatis                   ............................................................
Klebsiella oxytoca                        ............................................................

Burkholderiales bacterium                 PVKPIAQFNQRWLALLDKARQWPIHRGRMTIAVVGAGAGGVELVLSMQYRLRNELKALGR
Comamonadaceae bacterium                  PVKPIAHFNQRWLDLLERVRGL...RDRFTVAVVGGGAGGVELVLSMQYRLRQELQALGK
Hydrogenophaga flava                      PVKPIAHFNQRWLALLERVRSL...RDRFTVAVVGGGAGGVELVLSMQYRLRQELQALGK
Hydrogenophaga pseudoflava                PVKPIARFNQRWVNLLDRVKTH...AGKTTIAVVGGGAGGVELALAMQFRLRNELSTMGR
Limnobacter sp.                           PVKPIAHFNQRWLQLLDRVRGL...HSRFTIAVVGGGAGGVELVLSVQFRLRNELLKLGR


Methanothermococcus thermolithotrophicus  ............................................................
Methanocaldococcus fervens                ............................................................
Methanocaldococcus vulcanius              ............................................................
Methanocaldococcus jannaschii             ............................................................
Methanocaldococcus infernus               ............................................................

Cedecea davisae                           ............................................................
Escherichia fergusonii                    ............................................................
Citrobacter koseri                        ............................................................
Mannheimia granulomatis                   ............................................................
Klebsiella oxytoca                        ............................................................

Burkholderiales bacterium                 NPEDLQFVLLTAGETILPTHNPGVRARFARVLQARNVTVHTQTEVVEVSPGCLHTRDGRT
Comamonadaceae bacterium                  NPDLLQFVLLTAGETLLPTHNPRVRARFAHVLKERRVAVHTRAEVSQVSPGCLTTQDGRT
Hydrogenophaga flava                      SPDLLQFVLLTAGDSLLPTHNPRVRARFAHVLKERRVAVHTRAEVTQVSPGCLTTQDGRT
Hydrogenophaga pseudoflava                NPDELEFHLFTADADVLPTHNPGVRARFEKVLNERGVVLHRDAEVVQLDGHTLQTKQGER
Limnobacter sp.                           NPDLLRFVLLTAGDTILPTHNPGVRARFARVLKERHVAVHTRAEVTEVSPGCLHTQDGRT


                                                                                                      1
Methanothermococcus thermolithotrophicus  ........................................................DDAA
Methanocaldococcus fervens                ........................................................DDAA
Methanocaldococcus vulcanius              ........................................................DDAA
Methanocaldococcus jannaschii             ........................................................DDAS
Methanocaldococcus infernus               ........................................................DDAS

Cedecea davisae                           ........................................................DDAA
Escherichia fergusonii                    ........................................................DDAA
Citrobacter koseri                        ........................................................DDAA
Mannheimia granulomatis                   ........................................................DDAA
Klebsiella oxytoca                        ........................................................DDAA

Burkholderiales bacterium                 FDADDTLWVTQAGGPAWLKSTGLALDKQGFILVHPQLQTLNDPLVFAAGDIA....DDAA
Comamonadaceae bacterium                  FDADETMWVTQAGGPAWLQGTGLALDEHGFVCVNEYLQTPDDPKIYAAGDVASFLPDDAA
Hydrogenophaga flava                      FDADETMWVTQAGGPAWLQGTGLALDEHGFVCVNEYLQTPDDPKIFAAGDVASFLPDDAA
Hydrogenophaga pseudoflava                LHADEIMWVTQAGGAAWLKNTDLELDSRGFINVGPTLQTTRDAKIFAAGDIAN...DDAA
Limnobacter sp.                           FDADETLWVTQAGGPVWLQSTGLALDEHGFIQVNQQLQTLDDPKIFAVGDVA....DDAA
```

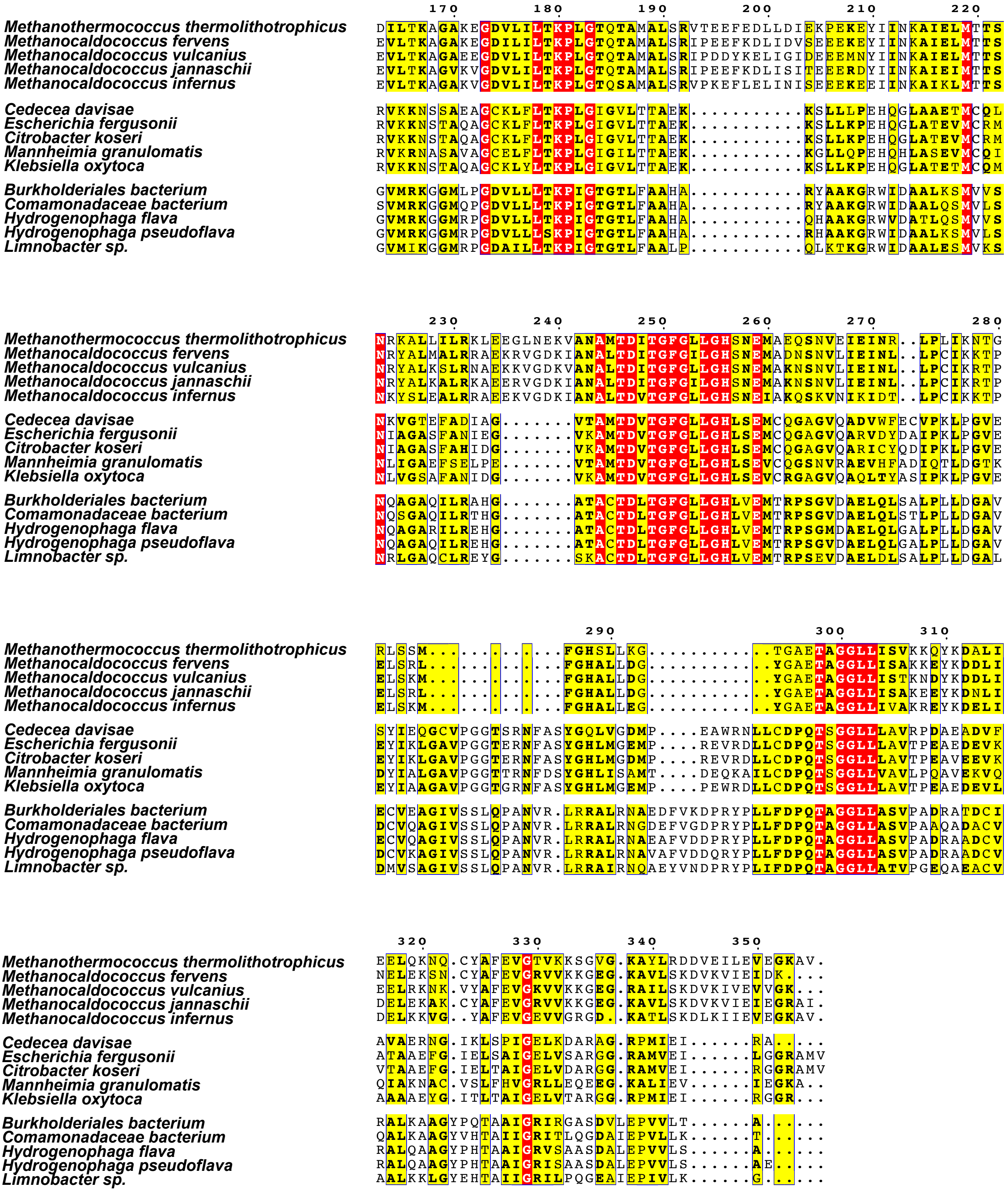
170 180 190 200 210 220
230 240 250 260 270 280
290 300 310
320 330 340 350
Methanothermococcus thermolithotrophicus
Methanocaldococcus fervens
Methanocaldococcus vulcanius
Methanocaldococcus jannaschii
Methanocaldococcus infernus
Cedecea davisae
Escherichia fergusonii
Citrobacter koseri
Mannheimia granulomatis
Klebsiella oxytoca
Burkholderiales bacterium
Comamonadaceae bacterium
Hydrogenophaga flava
Hydrogenophaga pseudoflava
Limnobacter sp.
